# All-in-One Dual CRISPR-Cas12a (AIOD-CRISPR) Assay: A Case for Rapid, Ultrasensitive and Visual Detection of Novel Coronavirus SARS-CoV-2 and HIV virus

**DOI:** 10.1101/2020.03.19.998724

**Authors:** Xiong Ding, Kun Yin, Ziyue Li, Changchun Liu

## Abstract

A recent outbreak of novel coronavirus (SARS-CoV-2), the causative agent of COVID-19, has spread rapidly all over the world. Human immunodeficiency virus (HIV) is another deadly virus and causes acquired immunodeficiency syndrome (AIDS). Rapid and early detection of these viruses will facilitate early intervention and reduce disease transmission risk. Here, we present an **A**ll-**I**n-**O**ne **D**ual **CRISPR**-Cas12a (termed “AIOD-CRISPR”) assay method for simple, rapid, ultrasensitive, one-pot, and visual detection of coronavirus SARS-CoV-2 and HIV virus. In our AIOD CRISPR assay, a pair of crRNAs was introduced to initiate dual CRISPR-Cas12a detection and improve detection sensitivity. The AIOD-CRISPR assay system was successfully utilized to detect nucleic acids (DNA and RNA) of SARS-CoV-2 and HIV with a sensitivity of few copies. Also, it was evaluated by detecting HIV-1 RNA extracted from human plasma samples, achieving a comparable sensitivity with real-time RT-PCR method. Thus, our method has a great potential for developing next-generation point-of-care molecular diagnostics.

## Introduction

SARS-CoV-2 (previously named 2019-nCoV) is a new coronavirus causing coronavirus disease 2019 (COVID-19) which first emerged in December 2019.^1^ As of 18 March 2020, based on the data provided by the World Health Organization,^2^ 191,127 people all over the world have been infection-confirmed and 7807 people have died. Human immunodeficiency virus (HIV) is one of another most dangerous virus and causes acquired immunodeficiency syndrome (AIDS). According to the World Health Organization (WHO), there were ∼37.9 million people living with HIV.^3^ Early detection of such pathogen infections during seroconversion window will facilitate early intervention (“test and treat”), which, in turn, may reduce disease transmission risk.

Polymerase chain reaction (PCR) method is the most commonly used technology for pathogen nucleic acid detection and has been considered as a “gold standard” for disease diagnostics due to high sensitivity and specificity.^4, 5, 6^ However, it typically relies on expensive equipment and well-trained personnel, all of which is not suitable for point of care diagnostic application. In recent decades, several isothermal amplification methods, such as recombinase polymerase amplification (RPA)^7^, loop-mediated isothermal amplification (LAMP)^8^, have been developed as attractive alternatives to conventional PCR method because of their simplicity, rapidity and low cost. However, there is still a challenge to apply it to develop a reliable POC diagnostics for clinical applications due to non-specific signals (e.g., false-positive).^9, 10^

Recently, RNA-guided CRISPR/Cas nuclease-based nucleic acid detection has shown great promise for the development of next-generation molecular diagnostics technology due to its high sensitivity, specificity and reliability.^11, 12^ For example, some Cas nucleases (e.g., Cas12a, Cas12b and Cas13a) perform strong collateral cleavage activities in which a crRNA-target-binding activated Cas can indiscriminately cleave surrounding non-target single-stranded nucleic acids.^13, 14, 15, 16, 17^ By combining with RPA preamplification, Cas13 and Cas12a have, respectively, been used to develop SHERLOCK (Specific High-sensitivity Enzymatic Reporter UnLOCKing) system^18^ and DETECTR (DNA Endonuclease-Targeted CRISPR Trans Reporter) system^14^ for highly sensitive nucleic acid detection. Apart from RPA method, some CRISPR-Cas-based nucleic acid sensors utilized the LAMP and PCR approaches, for instance, the CRISPR-Cas12b-assisted HOLMESv2 platform.^16^ However, these CRISPR-Cas-based detection methods typically require separate nucleic acid pre-amplification and multiple manual operations, which undoubtedly complicates the procedures and brings about contaminations.

In this study, we reported an **A**ll-**I**n-**O**ne **D**ual **CRISPR**-Cas12a (termed “AIOD-CRISPR”) assay for rapid, ultrasensitive, specific and visual detection of nucleic acid. Dual crRNAs are introduced to initiate highly efficient CRISPR-based nucleic acid detection. In our AIOD-CRISPR assay, all components for nucleic acid amplification and CRISPR detection are thoroughly mixed in a single, one-pot reaction system and incubated at a single temperature (e.g., 37 °C), eliminating the need for separate preamplification and amplified product transferring. As application examples, the AIOD-CRISPR assay was engineered to detect severe acute respiratory syndrome coronavirus 2 (SARS-CoV-2)^19^ and human immunodeficiency virus type 1 (HIV-1).^20^ Since SARS-CoV-2 and HIV-1 are retrovirus, we evaluated the performance of our AIOD-CRISPR assay by detecting both of their DNA and RNA. Especially, the test results of our AIOD-CRISPR assay can be directly visualized by naked eye. Therefore, we anticipate that the AIOD-CRISPR assay will facilitate CRISPR-based next-generation molecular diagnostics towards point-of-care applications.

## Results

### AIOD-CRISPR assay system

As shown in **Figure 1A**, the AIOD-CRISPR assay system used a pair of Cas12a-crRNA complexes generated by two individual crRNAs to bind two corresponding sites which are close to the recognition sites of primers in the target sequence. The Cas12a-crRNA complexes were separately prepared prior to being loaded into the solution containing two RPA primers, ssDNA-FQ reporters, recombinase, single-stranded DNA binding protein (SSB), strand-displacement DNA polymerase, and target sequences. When incubating the AIOD-CRISPR system in one pot at 37°C, RPA amplification is first initiated and exposes the binding sites of the Cas12a-crRNA complexes due to the strand displacement. Once the Cas12a-crRNA complexes bind the target sites, Cas12a endonuclease is activated and cleaves the nearby ssDNA-FQ reporters to produce fluorescence. Similarly, RPA amplified products are also the substrates to undertake the same process and continuously trigger CRISPR-Cas12a-based collateral cleavage activity. Previous study ^14, 17^ has demonstrated that this collateral cleavage activity is irrelevant to target strand cleavage in CRISPR-Cas12a system. Therefore, target sequences for the AIOD-CRISPR assay are not limited by the Cas12a’s protospacer adjacent motif (PAM) ^21^.

**Figure 1.**
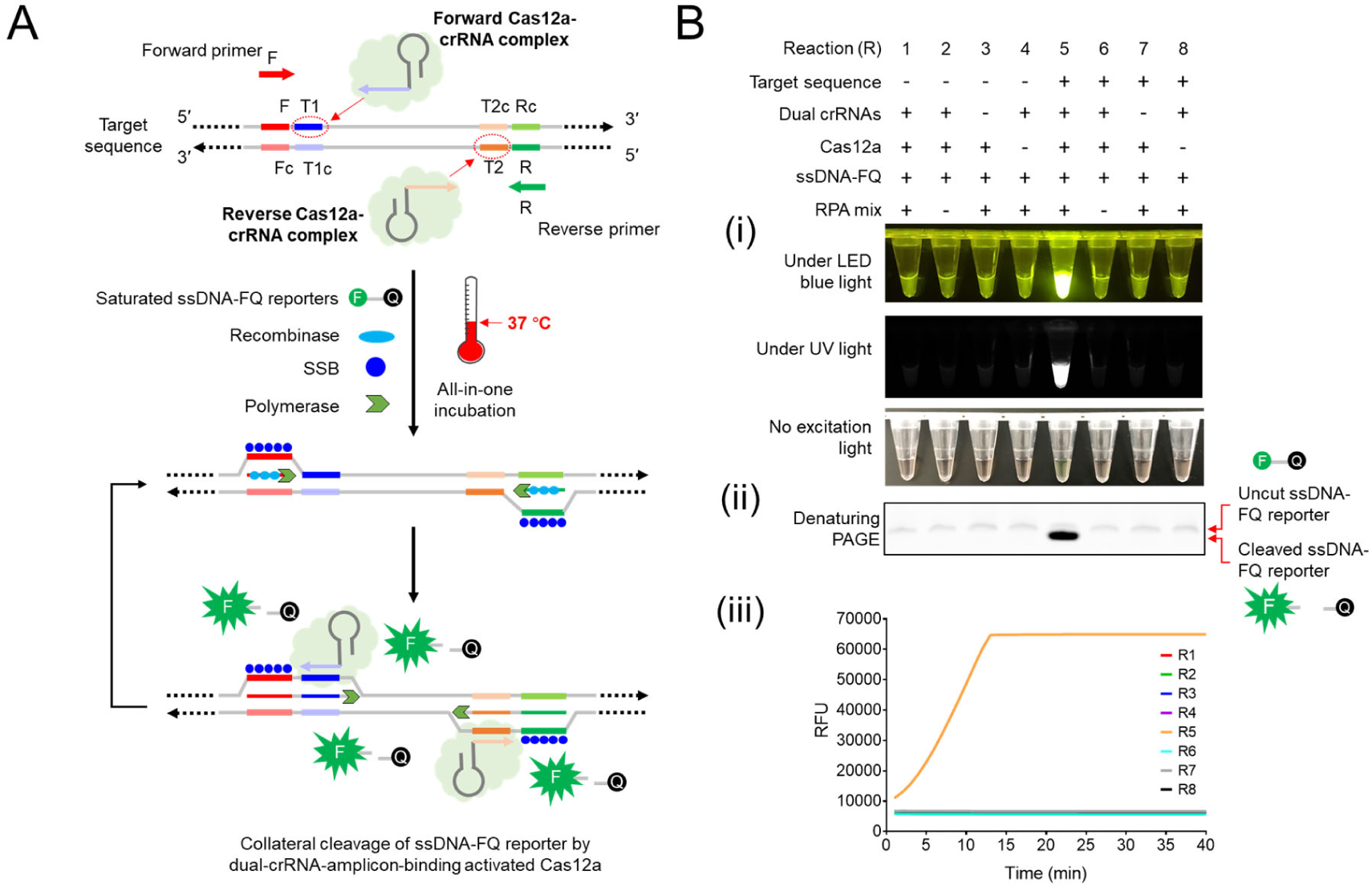
Design and working principle of the AIOD-CRISPR assay. (A) Schematic of the AIOD-CRISPR assay system. SSB, single-stranded DNA binding protein. (B) Development and evaluation of the AIOD-CRISPR assay system. The ssDNA-FQ reporter was labelled by 5’ 6-FAM (Fluorescein) fluorophore and 3’ Iowa Black^®^ FQ quencher. Recombinase polymerase amplification (RPA) mix from TwistAmp^®^ Liquid Basic kit was composed of 1× Reaction Buffer, 1× Basic E-mix, 1× Core Reaction Buffer, 14 mM MgOAc, 320 nM each of primers and 1.2 mM dNTPs. Dual crRNAs contained 0.64 μM each of crRNAs. HIV-1 p24 plasmid (1.2× 10^5^ copies), 8 μM of ssDNA-FQ reporters, and 0.64 μM EnGen^®^ Lba Cas12a (Cpf1) were used. (i) Eight reaction systems with various components and their endpoint images after 40 min incubation. (ii) Denaturing PAGE analysis of the AIOD-CRISPR products. (iii) Real-time fluorescence detection of the AIOD-CRISPR assay for eight reaction systems with various components.

To evaluate and develop our AIOD-CRISPR assay, we prepared and tested eight reaction systems (Reactions # 1-8) with various components (**Figure 1B (i)**). The ssDNA-FQ reporter was a 5 nt oligonucleotide (5’-TTATT-3’) labelled by 5’ 6-FAM (Fluorescein) fluorophore and 3’ Iowa Black^®^ FQ quencher. After incubated at 37°C for 40 min, only reaction # 5 containing the target sequence, dual crRNAs, Cas12a, and the RPA mix produced super-bright fluorescence signal, which could be directly visualized in a LED blue or UV light illuminator. Interestingly, even under ambient light environment without any excitation, a color change from orange-yellow to green was observed in the reaction tube # 5 by naked eyes. Furthermore, the assay products were verified by denaturing polyacrylamide gel electrophoresis (PAGE). As shown in **Figure 1B(ii)**, a strong band with shorter DNA size was observed only in the lane of the reaction # 5, which resulted from the cleaved ssDNA-FQ reporter with bright fluorescence. On the contrary, for other reaction systems, only weak bands with slightly longer DNA sizes were observed in their corresponding lanes due to fluorescence quench of the intact uncut ssDNA-FQ reporters. In addition, in real-time fluorescence detection, only the reaction # 5 showed an elevated fluorescence signals that saturated within 13 min (**Figure 1B(iii)**). Thus, these results show that our AIOD-CRISPR assay provides a simple, rapid, one-pot approach for target-specific nucleic acid detection.

Since RPA amplification reaction is initiated after adding MgOAc,^22^ we are interested in knowing if the nucleic acid amplification was initiated at room temperature during sample preparation in our AIOD-CRISPR assay system. We prepared two AIOD-CRISPR solutions (one positive and one negative) and allowed them to remain at room temperature for 10 min. As shown in **Figure S1**, no significant fluorescence change between positive and negative samples was observed in the AIOD-CRISPR solutions at room temperature. On the contrary, there was an obvious fluorescence increasing after just one-min incubation at 37°C (**Figure S1**). The fluorescence signal was eventually saturated at 15 min after incubated at 37°C. Therefore, our AIOD-CRISPR assay system is mainly triggered only after reaction temperature arrives at 37°C.

### Optimization of AIOD-CRISPR assay

Collateral cleavage of ssDNA-FQ reporter by the Cas12a is triggered by the binding of crRNA to target sites.^13, 14^ Thus, we hypothesized that higher binding opportunities could increase the collateral cleavage activity and eventually improve the detection sensitivity for AIOD-CRISPR. To prove this, we introduced a pair of crRNAs to recognize the corresponding target sites. A pUCIDT-AMP plasmid containing 300 bp HIV-1 p24 gene cDNA (p24 plasmid) was used as the target and three different designs for primers and crRNAs were investigated (**Figure 2A**). As shown in **Figure 2B**, AIOD-CRISPR with dual crRNAs (crRNA1+crRNA2) shows similar fluorescence curve as that with crRNA2, but was much better than using crRNA1 in real-time detection. Doubling the amount of crRNA1 or crRNA2 actually didn’t benefit the detection efficiency. On detection sensitivity, **Figure 2C** shows that AIOD-CRISPR with dual crRNAs was able to detect as low as 1.2 copies of the p24 plasmid templates within 40 min, while AIOD-CRISPR with single crRNA2 doesn’t detect it. Thus, introducing dual crRNAs is beneficial to improve the sensitivity of AIOD-CRISPR assay.

**Figure 2.**
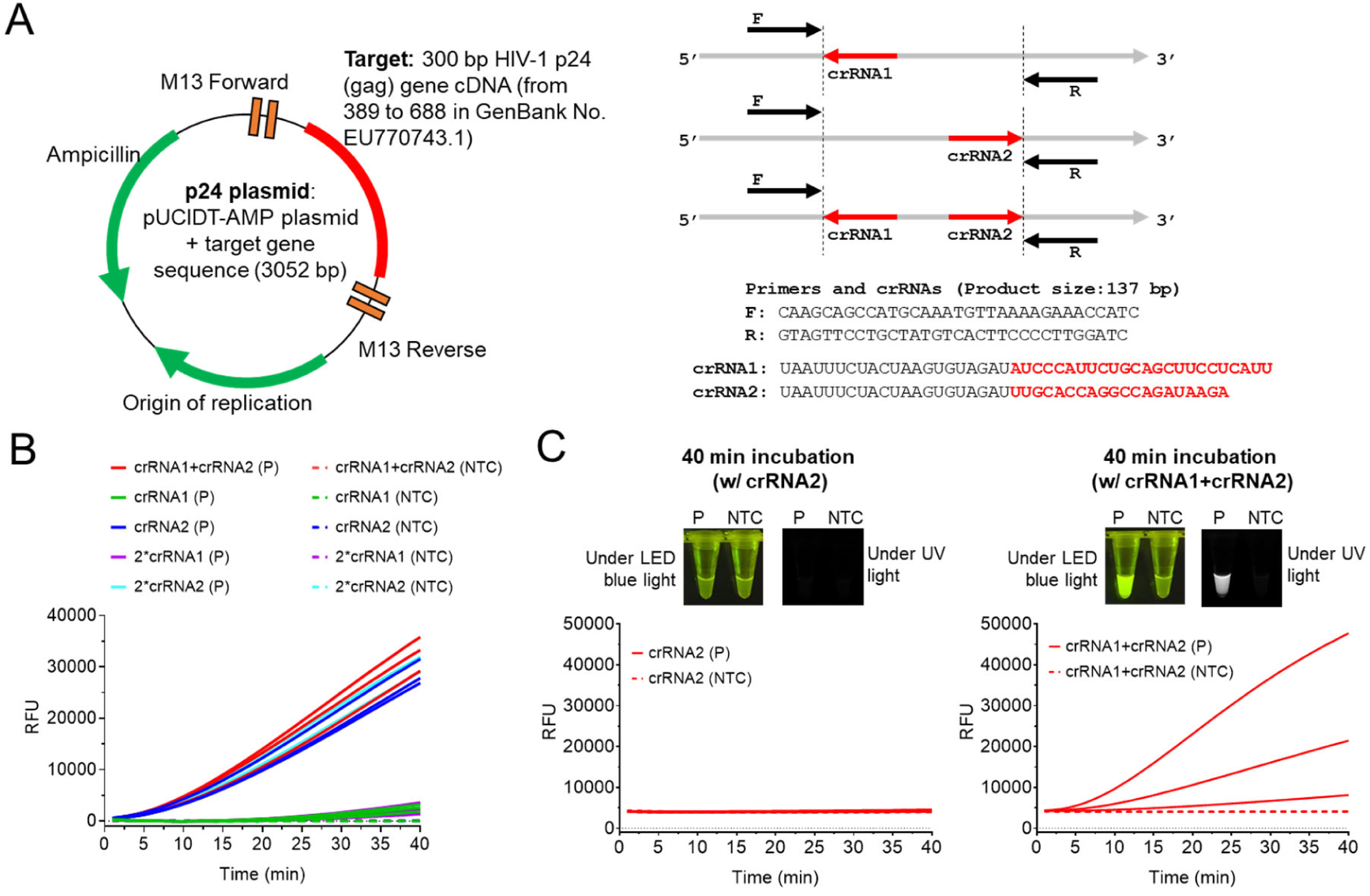
Comparison of the all-in-one CRISPR-Cas12a assay using dual crRNAs or single crRNA. (A) The pUCIDT-AMP plasmid containing 300 bp HIV-1 p24 gene cDNA (p24 plasmid) and the sequences of its primers and crRNAs (B) Real-time fluorescence detection of the all-in-one CRISPR-Cas12a assay using dual crRNAs (crRNA1+crRNA2) or single crRNA (crRNA1/crRNA2). 2*crRNA1/2*crRNA2 means doubling its amount. P, the positive reaction with 1.2× 10^3^ copies of HIV-1 p24 plasmids. (C) The CRISPR-Cas12a assays with dual crRNAs (crRNA1+crRNA2) or crRNA2 for the detection of 1.2 copies of HIV-1 p24 plasmids (P). Three replicates ran for each reaction with the plasmid. NTC, non-template control reaction. Each reaction contained 2 μM ssDNA-FQ. Fluorescence images were taken after 40 min incubation.

We further optimized ssDNA-FQ reporters in our the AIOD-CRISPR assay because the reporter concentration is crucial in fluorescence readout. As shown in **Figure S2A**, higher concentration of ssDNA-FQ reporters was used, shorter time was taken to reach saturated fluorescence intensity. As to threshold time and visual detection, the minimal concentration for saturated values was 4 μM (**Figure S2B-S2D**). Collateral cleavage by activated Cas12a is an activity of cutting nearby ssDNA-FQ reporters.^13, 14^ Thus, increasing ssDNA-FQ reporter concentration can strengthen the fluorescence signals. In addition, primer concentration also affected the AIOD-CRISPR’s efficiency. As shown in **Figure S3**, the optimal concentration should be 0.32 μM. Together, introducing dual crRNAs and using high concentration of ssDNA-FQ reporters enable efficient AIOD-CRISPR with 0.32 μM of each primer.

### HIV-1 detection by AIOD-CRISPR assay

To investigate its sensitivity for HIV-1 DNA detection, we first applied the optimized AIOD-CRISPR assay to detect various copies of HIV-1 p24 plasmid templates (from 1.2× 10^0^ to 1.2× 10^5^ copies). As shown in **Figure 3A**, the AIOD-CRISPR could detect as low as 1.2 copies HIV-1 p24 plasmid DNA in both real-time and endpoint visual detection in 40 min, which was further verified by the denaturing PAGE (**Figure 3B**). Although incubated for 40 min, the AIOD-CRISPR actually could identify 1.2 copies of HIV-1 DNA in just 1 min incubation based on endpoint fluorescence as shown in **Figure 3C**, which showed that our AIOD-CRISPR assay provides a super-fast and ultra-sensitive approach for nucleic acid detection.

**Figure 3.**
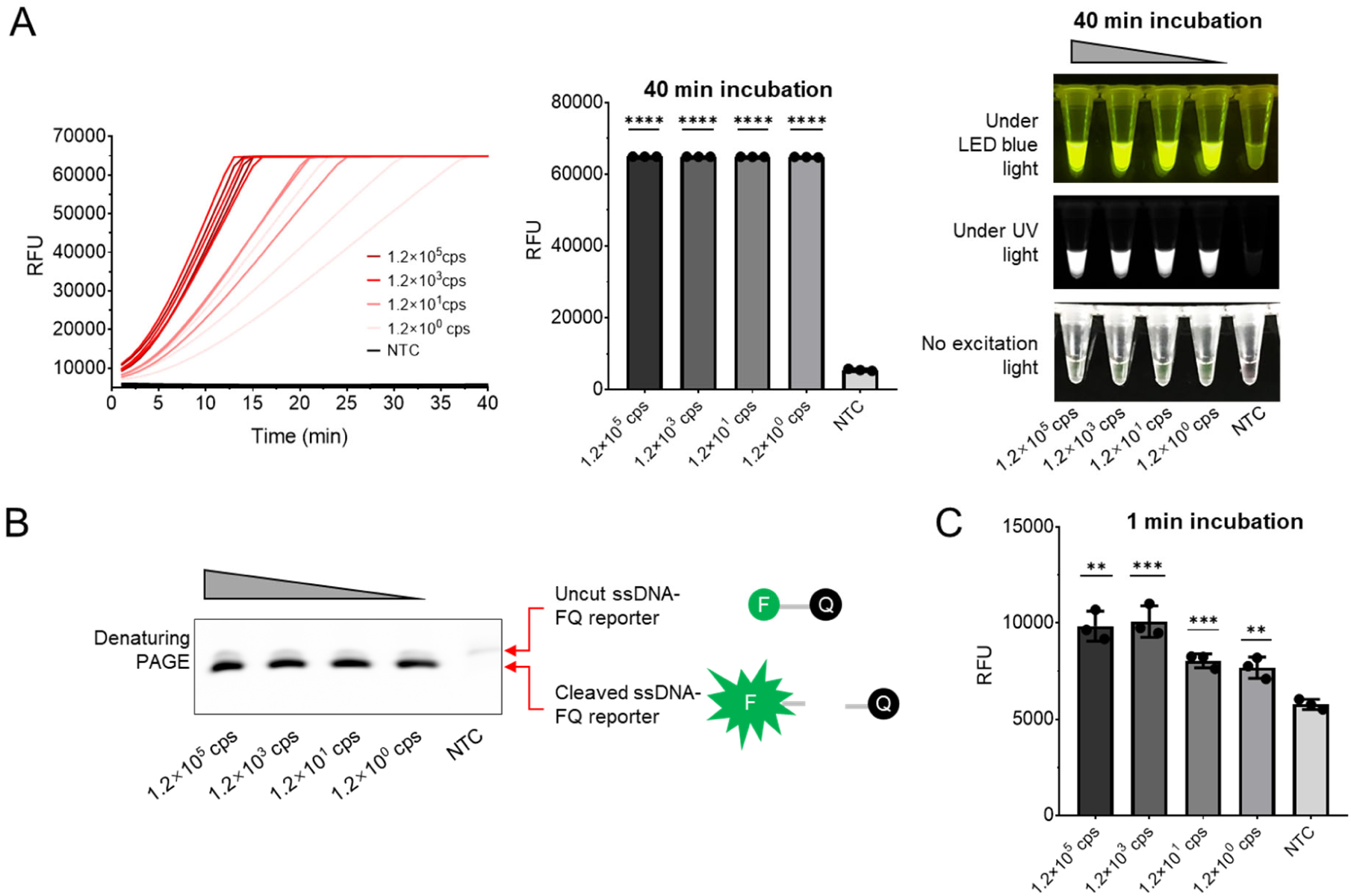
HIV-1 p24 plasmid (DNA) detection by the AIOD-CRISPR assay. (A) Real-time detection and endpoint fluorescence/visual detection. (B) Denaturing PAGE analysis of the AIOD-CRISPR products. (C) Endpoint fluorescence comparison after 1 min incubation at 37 °C. NTC, non-template control reaction. Three replicates ran for each reaction or test. Error bars represent the standard deviations at three replicates (n = 3). Unpaired two-tailed t-test was used to analyse the difference from NTC. \**P* < 0.05; \*\**P* < 0.01; \*\*\**P* < 0.001; \*\*\*\**P* < 0.0001. ns, not significant.

Next, we applied the AIOD-CRISPR assay to detect HIV-1 RNA sequence by adding Avian Myeloblastosis Virus (AMV) Reverse Transcriptase, namely reverse transcription AIOD-CRISPR (RT-AIOD-CRISPR). The HIV RNA target is a 1057-nt fragment of gag (p24 included) gene prepared using T7 promotor-tagged RT-PCR and T7 RNA polymerase-based transcription (**Figure S4A**). The detection region of AIOD-CRISPR was further verified by Sanger sequencing (**Figure S4B**). To achieve highly sensitive HIV RNA detection, we optimized the AMV concentration. As shown in **Figure S4C**, the RT-AIOD-CRISPR performed the highest efficiency with 0.32 U/μL AMV when incubated at 37°C. Furthermore, we investigated the sensitivity of our RT-AIOD-CRISPR assay using 1.1× 10^0^,1.1× 10^1^, 1.1× 10^3^, and 1.1× 10^5^ copies of HIV-1 gag RNA templates. **Figure 4A** shows that RT-AIOD-CRISPR incubated for 40 min at 37°C was able to consistently detect 11 copies RNA targets in real-time, endpoint fluorescence, and visual detections under LED blue and UV light. However, visual detection without excitation light only identified 1.1× 10^5^ copies RNA targets due to unsaturated fluorescence for other reactions. Additionally, the RT-AIOD-CRISPR’s sensitivity was further confirmed by denaturing PAGE analysis (**Figure S5**).

**Figure 4.**
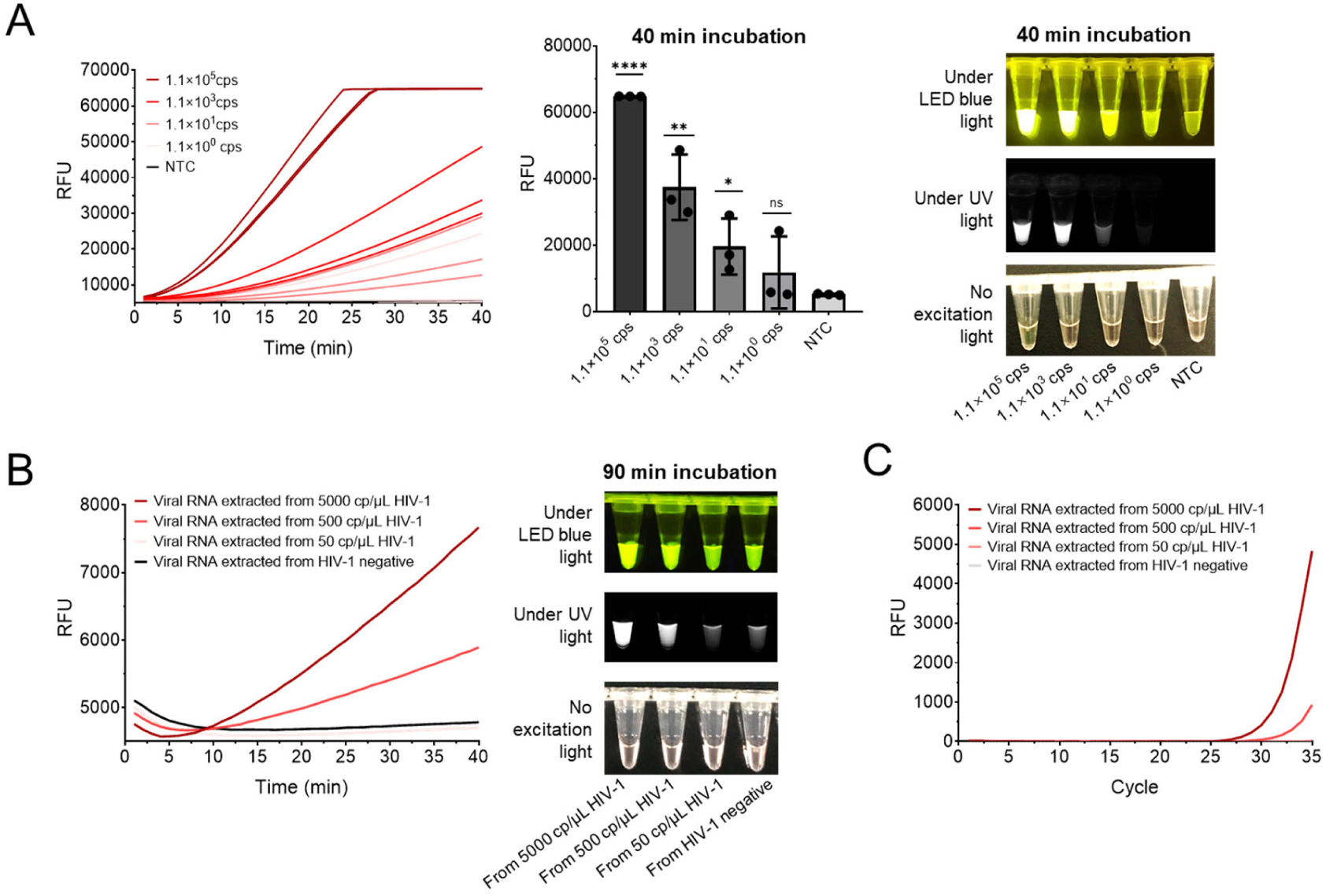
HIV-1 RNA detection by the RT-AIOD-CRISPR assay. (A) Sensitivity of real-time and endpoint fluorescence/visual detections. Three replicates were conducted for each test. NTC, non-template control reaction. Error bars represent the standard deviations at three replicates (n = 3). Unpaired two-tailed t-test was used to analyse the difference from NTC. \**P* < 0.05; \*\**P* < 0.01; \*\*\**P* < 0.001; \*\*\*\**P* < 0.0001. ns, not significant. (B) Real-time and visual AIOD-CRISPR assays for the detection of HIV-1 RNA extracted from human plasma samples. (C) Real-time RT-PCR assay to detect HIV-1 RNA extracted from human plasma samples. Negative, HIV-1 negative plasma samples.

In addition to assaying artificial HIV-1 gag RNA, we also analysed RT-AIOD-CRISPR’s performance using HIV-1 RNA extracted from plasma samples. As shown in **Figure 4B**, real-time RT-AIO-Cas12aR-based detection was able to detect the viral RNA extracted from 500 copies/μL HIV-1 plasma sample, though displaying low fluorescence change in less than 20 min. However, visual RT-AIOD-CRISPR detection took relatively long incubation time (up to 90 min) to achieve the similar sensitivity. The reduced efficiency of RT-AIOD-CRISPR likely results from some potential inhibitors or disruptors in the extracts. Despite this, real-time AIOD-CRISPR assay was still comparable to real-time RT-PCR assay regarding the same detection sensitivity (**Figure 4C and S6**).

### SARS-CoV-2 detection by AIOD-CRISPR assay

As shown in **Figure 5A**, a pUCIDT-AMP plasmid containing 384 nt SARS-CoV-2 N gene cDNA (N plasmid) was first prepared as the target to develop our AIOD-CRISPR assay. **Figure 5B** shows that our AIOD-CRISPR assay could detect 1.3 copies of SARS-CoV-2 N plasmids in both real-time and visual detections within 40 min, offering a rapid and nearly single-molecule level sensitive detection. To evaluate the detection specificity, we tested our AIOD-CRISPR assay using commercially available control plasmids containing the complete N gene from SARS-CoV-2 (SARS-CoV-2 _PC), SARS (SARS-CoV_PC), and MERS (MERS-CoV_PC), as well as the Hs_RPP30 control (Hs_RPP30_PC) with a portion of human RPP30 gene. **Figure 5C** shows that only the reaction with SARS-CoV-2_PC shows the positive signal in both real-time and visual detections, demonstrating that our developed AIOD-CRISPR assay possesses high specificity without cross reactions for non-SARS-CoV-2 targets.

**Figure 5.**
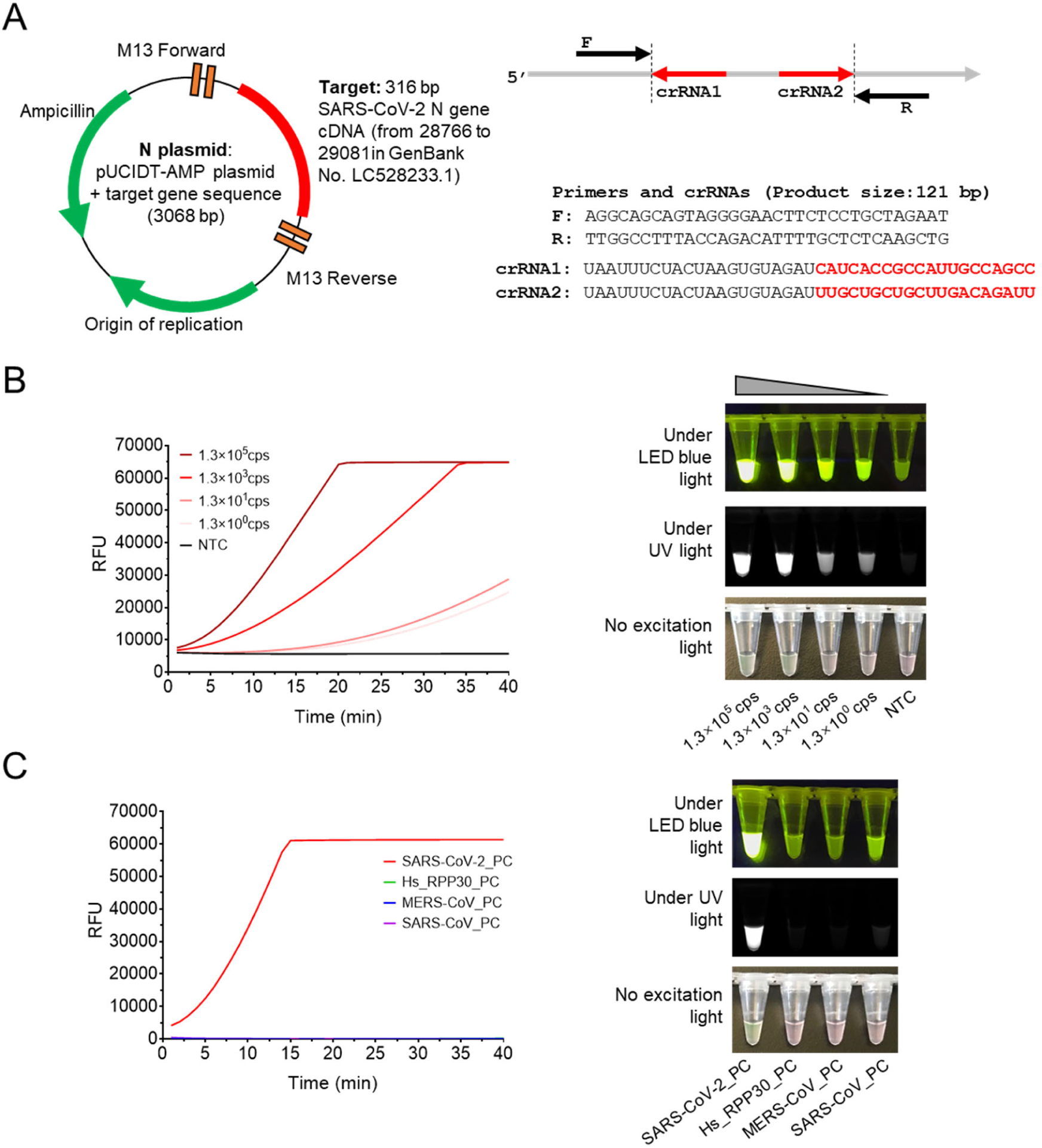
SARS-CoV-2 N DNA detection by the AIOD-CRISPR assay. (A) The pUCIDT-AMP plasmid containing 316 bp SARS-CoV-2 N gene cDNA (N plasmid) and the primers and crRNAs. (B) Real-time AIOD-CRISPR detection with various copies of SARS-CoV-2 N DNA. (C) Specificity assay of the AIOD-CRISPR assay on SARS-CoV-2 N detection. Tube images were taken after 40 min incubation.

Next, we used T7 promotor-tagged PCR and T7 RNA polymerase to prepare SARS-CoV-2 N gene RNA sequences to develop the RT-AIOD-CRISPR assay (**Figure 6A**). The detection region of RT-AIOD-CRISPR was verified by Sanger sequencing (**Figure 6B**). As shown in **Figure 6C**, both real-time and visual AIOD-CRISPR-based detections could detect down to 4.6 copies of SARS-CoV-2 N RNA targets in 40 min. Therefore, our AIOD-CRISPR assay is demonstrated to be a rapid, highly sensitive and specific SARS-CoV-2 detection method.

**Figure 6.**
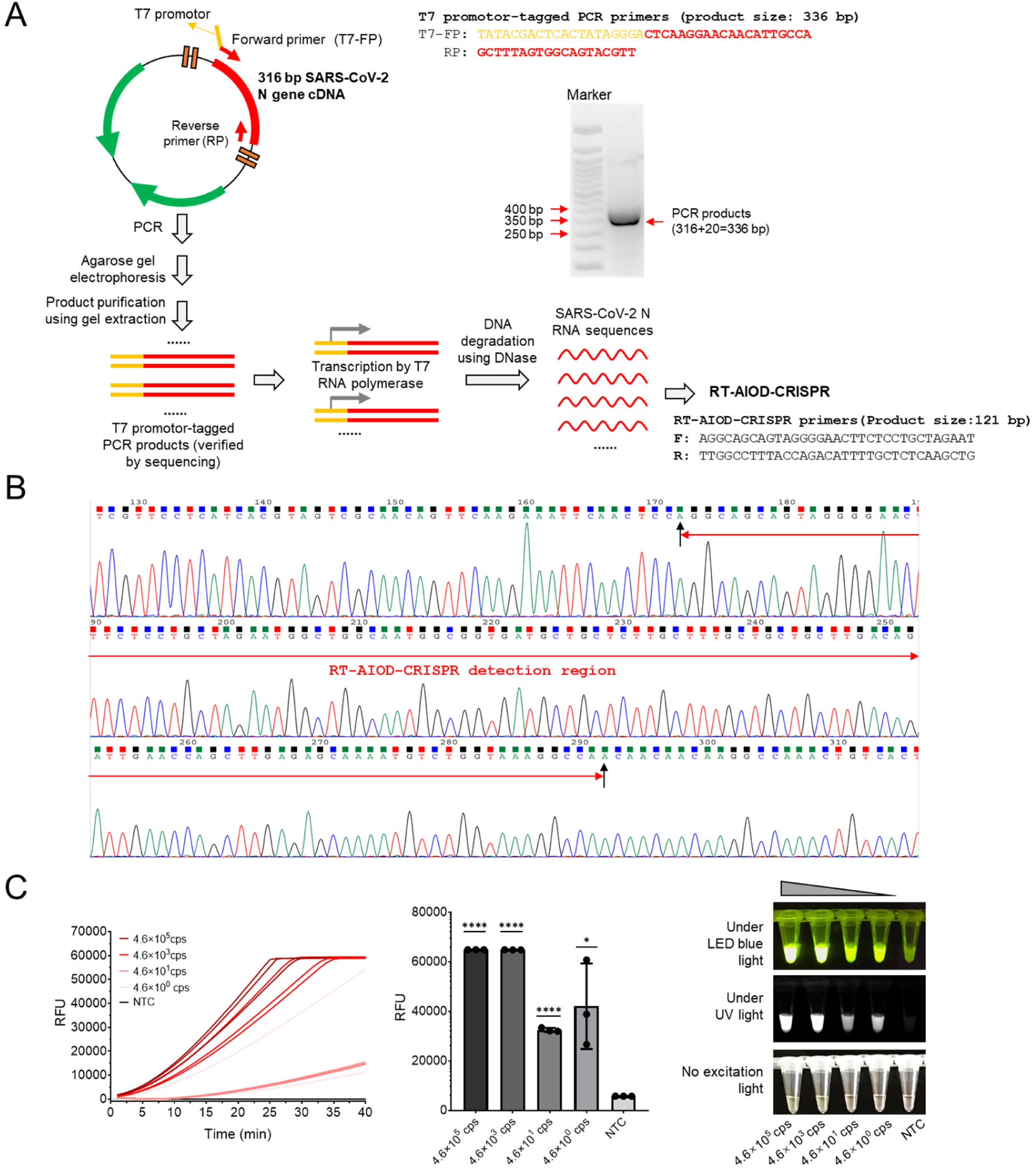
SARS-CoV-2 RNA detection by the RT-AIOD-CRISPR assay. (A) Protocol and PCR primers for preparing the SARS-CoV-2 N RNA sequences. (B) Sanger sequencing of the AIOD-CRISPR detection region in the prepared SARS-CoV-2 N RNA. (C) Sensitivity of real-time and endpoint fluorescence/visual AIOD-CRISPR detections. NTC, non-template control reaction. Three replicates were conducted for each test. Error bars represent the standard deviations at three replicates (n = 3). Unpaired two-tailed t-test was used to analyse the difference from NTC. \**P* < 0.05; \*\**P* < 0.01; \*\*\**P* < 0.001; \*\*\*\**P* < 0.0001. ns, not significant.

## Discussion

In this study, we describe a new CRISPR-Cas12a reaction, AIOD-CRISPR in which all components are incubated in a single reaction system to rapidly, highly sensitively and specifically detect nucleic acids without separate preamplification steps. Upon the design of AIOD-CRISPR, dual crRNAs are introduced to improve the sensitivity and high-concentration ssDNA-FQ reporters are added to strengthen the detection signals. In addition to real-time detection, AIOD-CRISPR assay is also beneficial for point-of-care detection because its detection results can be visually judged based on the fluorescence and color change of reaction solutions. As proof-of-concept assays, AIOD-CRISPR has been successfully developed for SARS-CoV-2 and HIV-1 detection.

Compared to previously reported CRISPR-based nucleic acid detection,^14, 16, 18, 23, 24, 25^ our versatile and robust AIOD-CRISPR has some distinctive advantages. First, AIOD-CRISPR system is a true single reaction system. All the components are prepared prior to incubation, thoroughly circumventing the preamplification of target nucleic acids,^14^ physical separation of Cas enzyme,^25^ or the requirement of specially designed tubes or devices^26^. Second, AIOD-CRISPR is a true isothermal nucleic acid detection method. AIOD-CRISPR is conducted in one-step and one-pot format at one temperature, eliminating the need of expensive thermocycler in PCR methods and the initial denaturing of dsDNA targets in isothermal nucleic acid amplification techniques such as LAMP ^8^ and SDA ^27^. Third, AIOD-CRISPR-based detection is very fast, robust, highly specific, and nearly single-molecule sensitive. In the application examples of HIV-1 and SARS-CoV-2 detections, our engineered AIOD-CRISPR without preamplification is able to detect as low as 1.2 copies DNA targets and 4.6 copies RNA targets in 40 min incubation. Especially in detecting HIV-1 p24 plasmids, AIOD-CRISPR can stably identify 1.2 copies in just 1 min incubation. For emerging SARS-CoV-2, in addition to high sensitivity, the AIOD-CRISPR also performs high detection specificity. Compared to the reported real-time RPA ^28^, AIOD-CRISPR shows a robust HIV-1 detection with very low background. Fourth, the AIOD-CRISPR enables one-step CRISPR-Cas12a-based RNA detection. By supplementing AMV reverse transcriptase, AIOD-CRISPR can be easily developed as one-step RT-AIOD-CRISPR to detect RNA targets such as HIV-1 and SARS-CoV-2 RNAs, which facilitates the CRISPR-Cas12a-based RNA detection without the preparation of cDNA. On detecting HIV-1 plasma samples, AIOD-CRISPR is comparable to real-time RT-PCR assay.

In summary, AIOD-CRISPR using a true single reaction system is a rapid, robust, highly specific, and nearly single-molecule sensitive isothermal nucleic acid detection method. This straightforward CRISPR-Cas12a nucleic acid sensing system has great potential in enabling CRISPR-based next-generation molecular diagnostics towards point-of-care, quantitation, and digital analysis.

## Methods

### Materials

Oligonucleotides (primers), ssDNA-FQ reporters, pUCIDT (Amp) plasmids with customized gene sequences, and the control plasmids containing the complete N gene from SARS-CoV-2, SARS, and MERS, as well as the Hs_RPP30 control were synthesized by or purchased from Integrated DNA Technologies (Coralville, IA). AcroMetrix™ HIV-1 plasma control samples, dNTP Set (100 mM), Gel Loading Buffer II (Denaturing PAGE), PureLink™ Quick Gel Extraction Kit and TURBO DNA-free™ Kit were purchased from Thermo Fisher Scientific (Waltham, MA). EvaGreen^®^ dye (20×) was purchased from Biotium (Fremont, CA). TEMED, (NH_4_)_2_S_2_O_8_, 30% acrylamide/bis-acrylamide solution, 10× TBE Buffer, and SsoAdvanced™ Universal SYBR^®^ Green PCR Supermix were purchased from Bio-Rad Laboratories (Hercules, CA). EnGen^®^ Lba Cas12a (Cpf1) (100 µM), deoxynucleotide (dNTP) mix (10 mM of each), and Avian Myeloblastosis Virus (AMV) Reverse Transcriptase (10,000 units/mL) were purchased from New England BioLabs (Ipswich, MA). QIAamp^®^ Viral RNA Mini Kit, RNeasy^®^ MinElute™ Cleanup Kit, and QIAGEN^®^ OneStep RT-PCR Kit were purchased from QIAGEN (Frederick, MD). RiboMAX™ Large Scale RNA Production Systems-T7 was purchased from Sigma-Aldrich (St. Louis, MO). TwistAmp^®^ Liquid Basic Kit was purchased from TwistDx™ Limited (Maidenhead, UK). The LED blue light illuminator (Maestrogen UltraSlim) was purchased from Fisher Scientific (Pittsburgh, PA).

### AIOD-CRISPR assays

The AIOD-CRISPR system was prepared separately as Component A, B and C. Component A contained 1× Reaction Buffer, 1× Basic E-mix, 14 mM MgOAc, 320 nM each of primers, and 1.2 mM dNTPs. Component B consisted of 4 μM of ssDNA-FQ reporters and 1× Core Reaction Buffer. Component C was the Cas12a-crRNA mix with 0.64 μM each of crRNAs and 0.64 μM EnGen^®^ Lba Cas12a. The concentration in each component was calculated based on the finally assembled 25-μL AIOD-CRISPR system. In a typical AIOD-CRISPR assay, 1 μL of the target solution was mixed with 20 μL of Component A and 2.5 μL of Component B. This assembled mixture was then mixed with 1.5 μL of Component C to form final 25 μL of AIOD-CRISPR system. For RT-AIOD-CRISPR assays, most components were the same as those in the AIOD-CRISPR system above, except supplementing 0.32 U/μL AMV Reverse Transcriptase in Component A. Real-time fluorescence detection was carried out in the Bio-Rad CFX96 Touch™ Real-Time PCR Detection System. Visual detection was accomplished through imaging the tubes in the LED blue light illuminator or the Bio-Rad ChemiDoc™ MP Imaging System with its built-in UV channel. For visual detection based on the reaction solution’s color change, 8 μM of ssDNA-FQ reporters should be used. All the reactions were incubated at 37°C for 40 min or the denoted time in figures. The endpoint fluorescence was the raw fluorescence determined by the Real-Time PCR Detection System. The unpaired t-test was applied for the statistical analysis in GraphPad Software Prism 8. A saturated fluorescence intensity was the maximum intensity which the Real-Time PCR Detection System could determine. After reaction, the AIOD-CRISPR solution was mixed with isometric Gel Loading Buffer II prior to 15% denaturing PAGE with 8 M urea and gel imaging in the Imaging System.

### In vitro RNA preparation using T7 RNA polymerase

For HIV-1 gag RNA, OneStep RT-PCR composed of 1× QIAGEN OneStep RT-PCR Buffer, 0.4 mM of each dNTP, 0.6 μM of each primer, 2.0 μL of QIAGEN OneStep RT-PCR Enzyme Mix, and 1.0 μL of the viral RNA extracted from HIV-1 plasma control was conducted to amplify the 1057 nt gag sequence. The thermal cycling protocol included 30 min at 50°C for reverse transcription, 15 min at 95°C for initial PCR activation, 35 cycles of the 3-step cycling (1 min at 94°C for denaturation, 1 min at 55°C for annealing, and 1 min at 72°C for extension), and 10 min at 50°C for final extension. For SARS-CoV-2 N RNA, the PCR system contained 1× SsoAdvanced™ Universal SYBR^®^ Green PCR Supermix, 0.4 μM of each primer, and 1.0 μL of 1.2× 10^5^ copies/μL HIV-1 p24 plasmid solution was used to amplify the 316 nt N sequence. The thermal cycling was 2.5 min at 98°C for initial denaturation, 35 cycles of 15 s at 95°C for denaturation and 30 s at 60°C for annealing and extension. The products of PCR/RT-PCR were all confirmed by agarose gel electrophoresis and Sanger sequencing. Afterwards, the products with the accurate sizes were extracted and purified using the Gel Extraction Kit. In vitro transcription was achieved through incubating the reaction system containing 8 μL of 5× T7 Transcription Buffer, 3 μL each of 100 mM rNTPs, 4 μL of the Enzyme Mix with T7 RNA polymerase, 16 μL of the gel-extracted PCR/RT-PCR products at 37°C for 4 h. Then, the transcription products were treated by DNase (from the TURBO DNA-free TM Kit) to degrading the DNA and the RNA was extracted and purified using the RNeasy^@^ MinElute™ Cleanup Kit. The purity and concentration of the collected RNA were determined using NanoDrop™ One/One^C^ Microvolume UV-Vis Spectrophotometry (Thermo Fisher Scientific).

### OneStep RT-PCR assay

QIAGEN^®^ OneStep RT-PCR Kit was used for the RT-PCR assay. The primers (FP: 5’-ATTATCAGAAGGAGCCACC-3’; RP: 5’-CATCCTATTTGTTCCTGAAGG-3’) for HIV-1 RNA detection were obtained from the reported literature.^29^ According to the instruction manual, the OneStep RT-PCR assay (50 μL) contained 1× QIAGEN OneStep RT-PCR Buffer, 400 µM of each dNTP, 600 nM each of primers (FP and RP), 2.0 μL of QIAGEN OneStep RT-PCR Enzyme Mix, 0.8× EvaGreen^®^ dye, and 5.0 μL of the RNA template solution. The thermal cycling protocol included 30 min at 50°C for reverse transcription, 15 min at 95°C for initial PCR activation step, 35 cycles of the 3-step cycling (30 s at 94°C for denaturation, 30 s at 55°C for annealing, and 1 min at 72 °C for extension), and 10 min at 72°C for final extension, followed by the melt-curve analysis (from 65 °C to 95 °C with 0.5 °C increment). Real-time OneStep RT-PCR assay was conducted in the CFX96 Touch™ Real-Time PCR Detection System and the plate read was set at the annealing in the 3-step cycling.

## Acknowledgements

The work was supported, in part, by R01EB023607, R01CA214072, and R21TW010625.

## Conflict of Interest Disclosures

University of Connecticut has filed a patent application on the methods described, and Changchun Liu and Xiong Ding are named as inventors.

## Supporting Information

**Figure S1.**
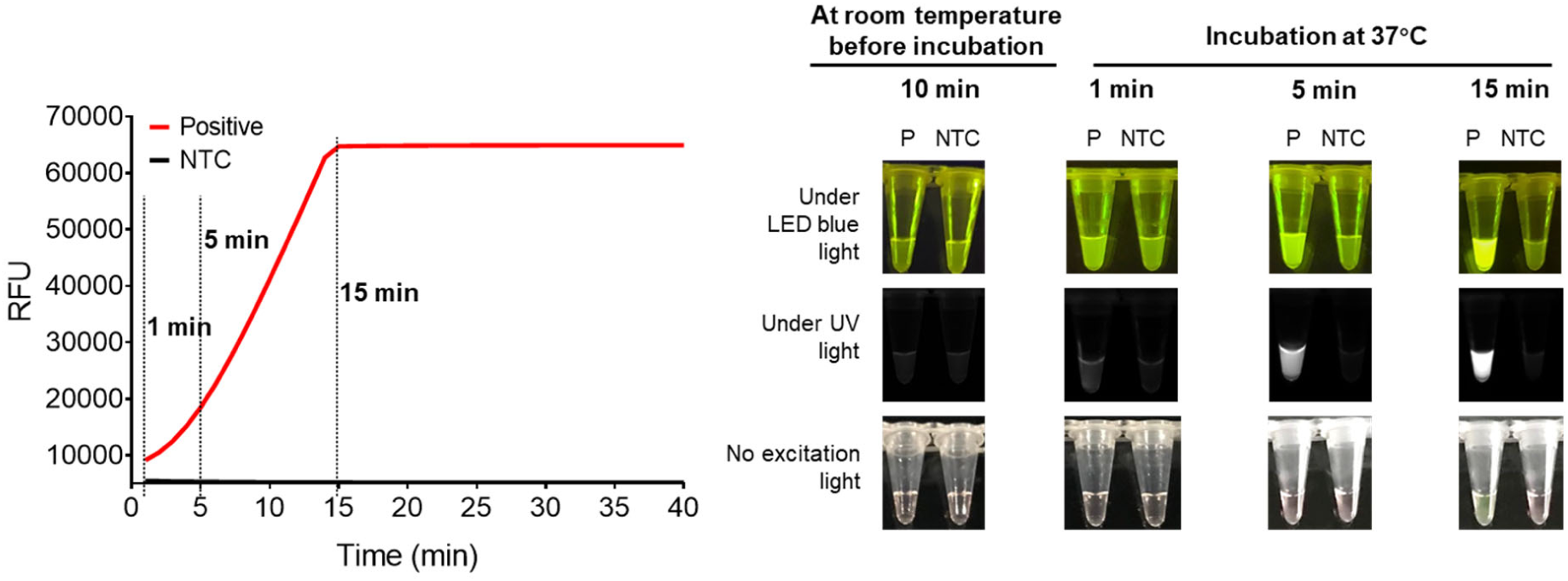
The AIOD-CRISPR assay at room temperature for 10 min before incubation and incubated at 37°C for 1, 5, and 15 min. Positive (P), the reaction with 1.2× 10^5^ copies of HIV-1 p24 plasmids. NTC, non-target control reaction.

**Figure S2.**
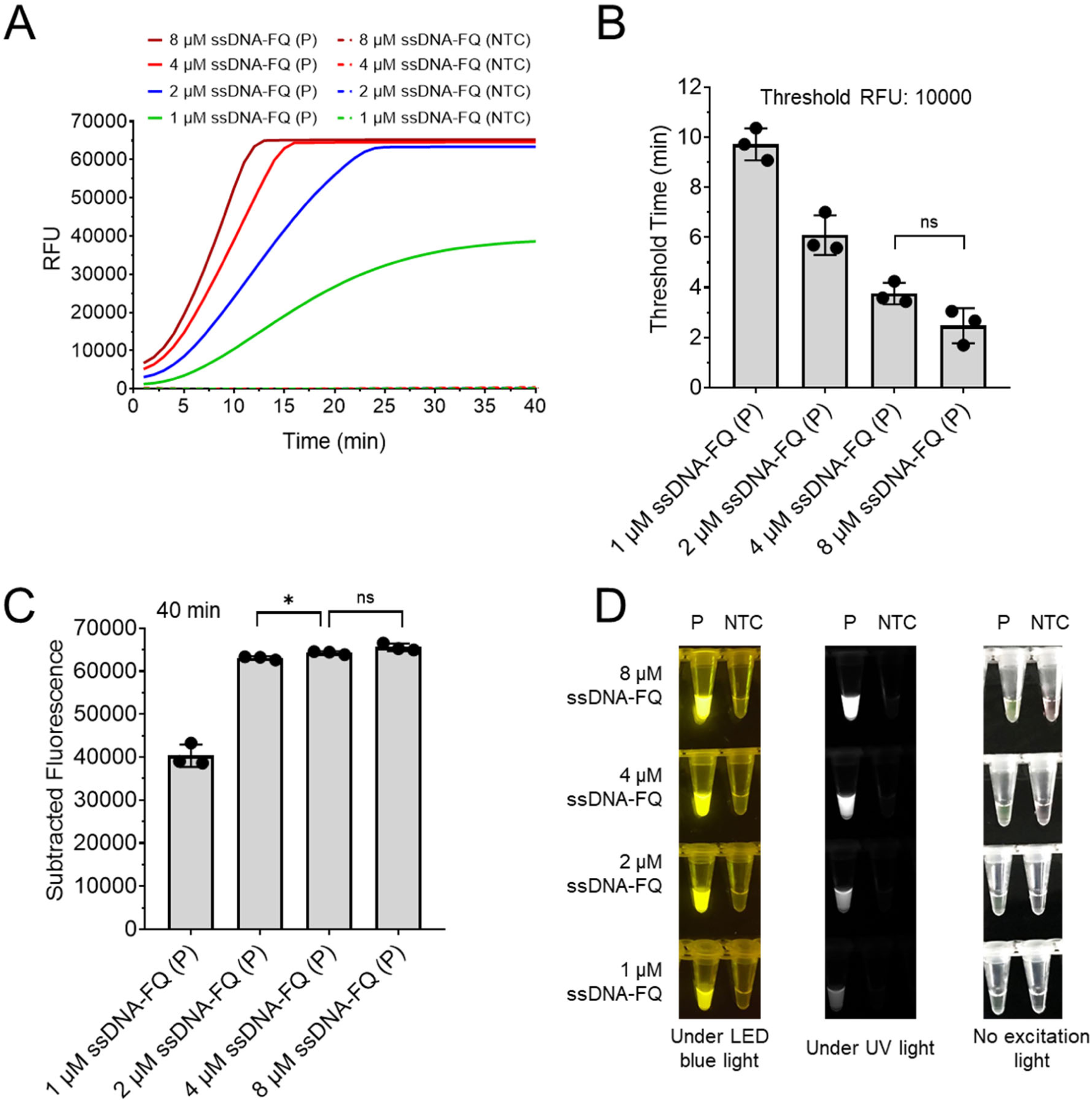
The AIOD-CRISPR assay with various concentrations of ssDNA-FQ reporters. (A) Real-time fluorescence detection. (B) Threshold time comparison. (C) Background-subtracted fluorescence comparison after 40 min incubation. (D) Visual detection comparison after 40 min incubation. P, the positive reaction with 2.4× 10^5^ copies of HIV-1 p24 plasmids. NTC, non-target control reaction. Error bars represent the standard deviations at three replicates (n = 3). Unpaired two-tailed t-test was used to analyse the difference. *P < 0.05; **P < 0.01; ***P < 0.001; ****P < 0.0001. ns, not significant.

**Figure S3.**
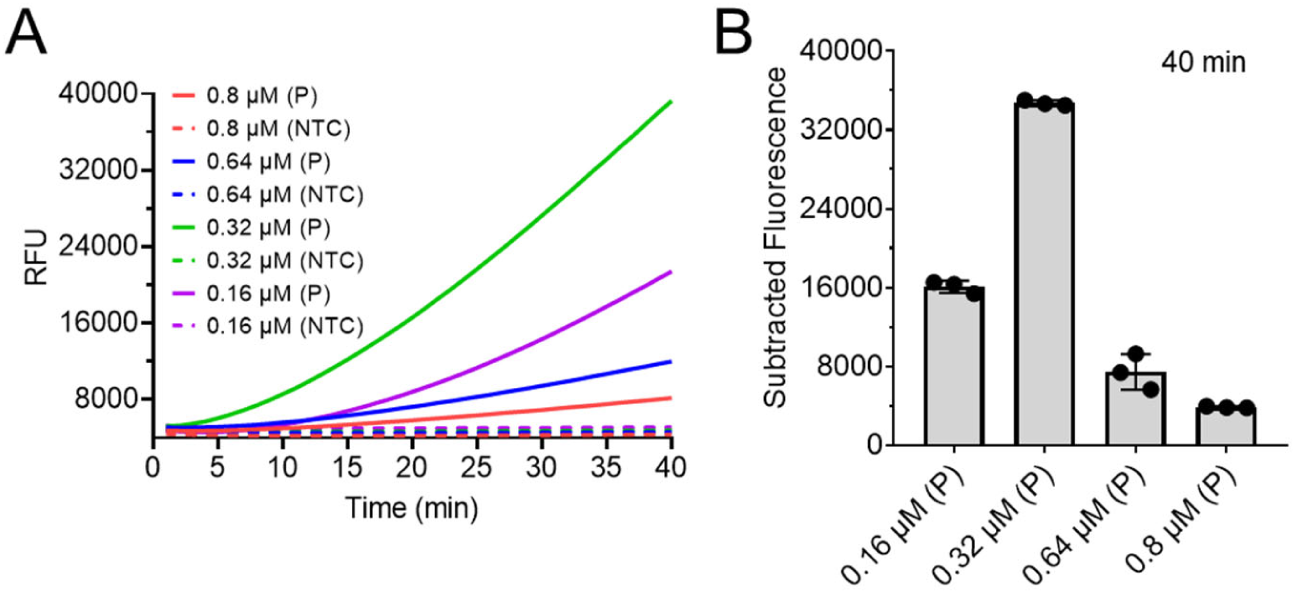
The AIOD-CRISPR assay with various concentrations of primers. (A) Real-time fluorescence detection. (C) Background-subtracted fluorescence comparison after 40 min incubation. P, the positive reaction with 1.2× 10^3^ copies of HIV-1 p24 plasmids. NTC, non-target control reaction. Error bars represent the standard deviations at three replicates (n = 3). Each reaction contained 2 μM ssDNA-FQ.

**Figure S4.**
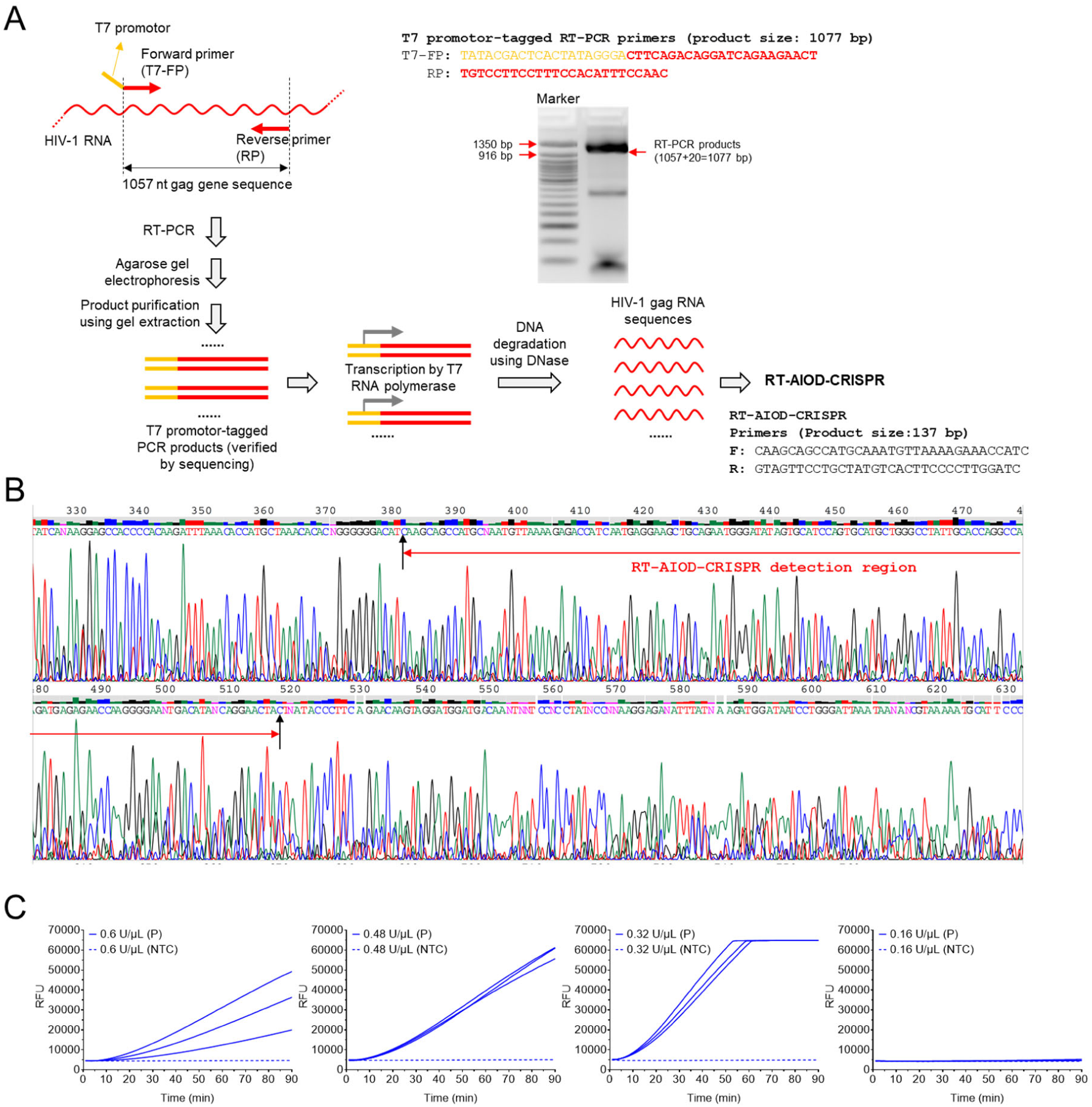
Development of RT-AIOD-CRISPR assay. (A) Protocol and RT-PCR primers for preparing HIV-1 gag RNA sequences. (B) Sanger sequencing of RT-AIO-Cas12aR-based detection region in the prepared HIV-1 gag RNA. (C) The RT-AIOD-CRISPR assay using various concentrations of AMV reverse transcriptase. Three replicates ran for each reaction or test. P, the positive reaction with 1.1× 10^3^ copies of HIV-1 gag RNA sequences. NTC, non-target control reaction.

**Figure S5.**
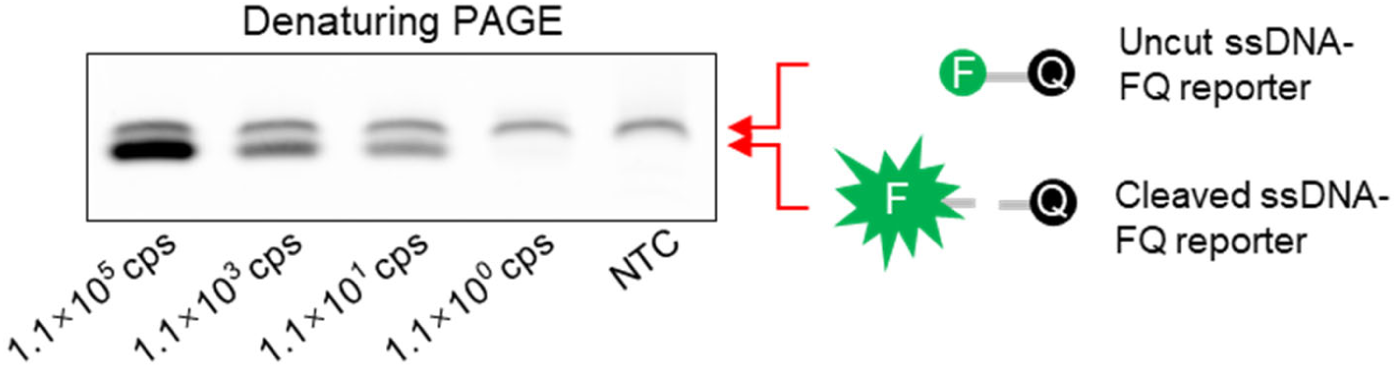
Denaturing PAGE analysis of RT-AIOD-CRISPR with various copies of HIV-1 gag RNA sequences.

**Figure S6.**
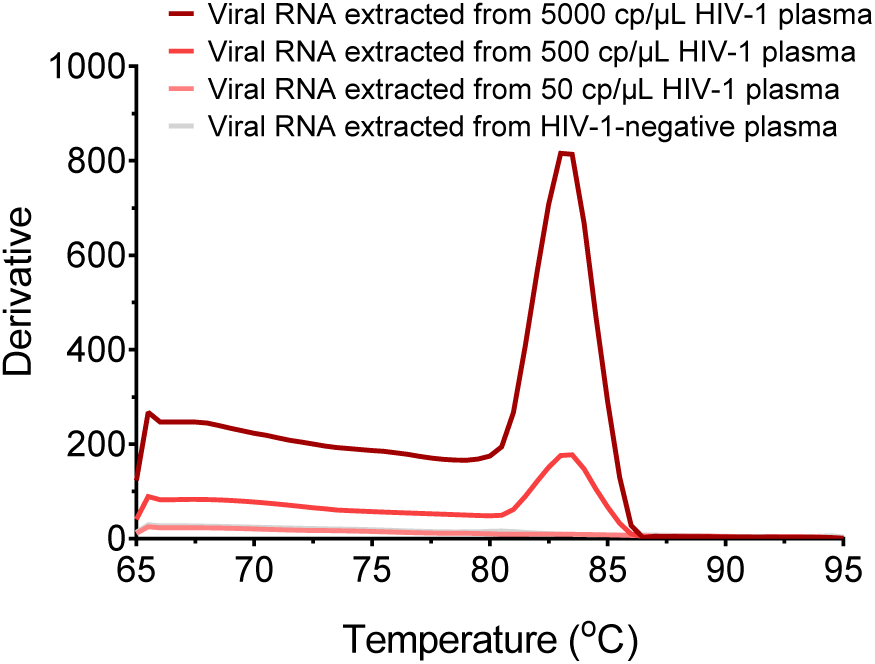
The melting curves for the OneStep RT-PCR products on detecting viral RNA extracted from various copies per microliter of HIV-1 plasma. The melting temperature of the products was about 83.5 °C.

